# Mapping movement, mood, motivation, and mentation in the subthalamic nucleus

**DOI:** 10.1101/168302

**Authors:** Amritha Gourisankar, Sarah A. Eisenstein, Nicholas T. Trapp, Jonathan M. Koller, Meghan C. Campbell, Mwiza Ushe, Joel S. Perlmutter, Tamara Hershey, Kevin J. Black

## Abstract

The anatomical connections of the subthalamic nucleus (STN) have driven hypotheses about its functional anatomy, including the hypothesis that the precise anatomical location of STN deep brain stimulation (DBS) determines the variability of motor and non-motor responses across Parkinson disease (PD) patients. We previously tested that hypothesis using a three-dimensional (3D) statistical method to interpret the acute effects of unilateral DBS at each patient’s clinically optimized DBS settings and active contact. Here we report a similar analysis from a new study in which DBS parameters were standardized and DBS locations were chosen blind to clinical response. In 74 individuals with PD and STN DBS, STN contacts were selected near the dorsal and ventral border of the STN contralateral to the more affected side of the body. Participants were tested off PD medications in each of 3 conditions (ventral STN DBS, dorsal STN DBS and DBS off) for acute effects on mood, apathy, working memory, response inhibition and motor function. Voltage, frequency, and pulse width were standardized, and participants and raters were blind to condition. In a categorical analysis, both dorsal and ventral STN DBS improved mean motor function without affecting cognitive measures. Dorsal STN DBS induced greater improvement in rigidity than ventral STN DBS, whereas ventral STN DBS was more effective for improving anxiety and mood. In the 3D analysis, contact location was significant only for bradykinesia and resting tremor, with the greatest improvement occurring with DBS in dorsal STN and zona incerta. These results provide new, direct functional evidence for the anatomically-derived model of STN using the novel 3D analysis, in which motor function is most represented in dorsal STN. However, our data suggest that functional segregation between motor and non-motor areas of the STN is limited, since locations that induced improvements in motor function and mood overlapped substantially.

## Introduction

Parkinson disease (PD) is the second most common neurodegenerative disease.^1^ PD varies in its presentation; symptoms may include disturbed sleep, depressive symptoms, apathy and cognitive complications in addition to classic motor features such as bradykinesia, rigidity and tremor.^2^ Deep brain stimulation of the subthalamic nucleus (STN DBS) can improve many of the motor symptoms,^3^ but changes in mood, motivation and cognition also occur and may be either beneficial or detrimental to the patient.^4^ In fact, clinical results vary substantially among patients. Some evidence suggests that the location of stimulation within or around the STN may contribute to the motor, mood, and cognitive effects of STN-DBS, given its relatively segregated anatomical connections to motor, somatosensory, and limbic neural circuits.^5^ However, the methods used to test this hypothesis in the past have had limitations including not examining the entire relevant volume of the brain^6,7,8^, not determining the statistical significance of relationships between behavior and DBS site^9,10,11,12^, or not correcting for Type 1 errors due to the multiple comparisons inherent in 3D statistical maps with many data points (i.e. voxels)^13^.

Some studies examined the effects of DBS on neuronal response with reference to the volume of tissue predicted to be activated based on electrical field models.^14^ We combined the anatomical location of the stimulated electrode with clinical data to produce statistical images that demonstrate DBS locations associated with improvement and worsening of each measured symptom, and determined overall statistical significance from these images using a permutation approach.^15^ This method avoids the issues noted above, and identifies whether location relates to clinical response in a statistically rigorous manner controlled for multiple comparisons.

Using this method, we previously examined the acute effects of unilateral STN DBS in PD, using each person’s clinically optimized stimulation parameters and electrode contacts. Mood, cognition and motor function were assessed with DBS OFF and ON at least 8 hours after the most recent dose of PD-related medication. The 3D analyses suggested that location of stimulation was significantly associated with mood, cognition, and some motor outcomes.^15^ Most motor measures improved with DBS everywhere in the STN, while a few motor, cognitive, and mood measures differed depending on the location of stimulation. An important weakness of that study was that stimulation parameters (e.g. voltage) differed across individuals, which could differentially impact behavior. The stimulation parameters used and the contact chosen were determined through the clinical programming process, so the results could not distinguish whether all participants would have had similar motor benefit with DBS anywhere in the STN, or whether the ideal DBS location simply varied by participant. Therefore, in this new study, all PD participants had separate, blinded, unilateral stimulation conditions at both dorsal *and* ventral STN locations, chosen by brain imaging blind to clinical results. All stimulation parameters were maintained across condition and participant. We hypothesized that our findings would be qualitatively similar to those in our previous report, but that effects might be more striking due to the consistent stimulation parameters and the more uniform approach to selecting DBS locations in both dorsal and ventral STN.

## Materials and methods

### Participants

Seventy-four PD patients were recruited through the Movement Disorders Center at Washington University St. Louis School of Medicine (WUSM). Inclusion criteria included bilateral STN DBS therapy for clinically definite PD, as previously defined^16^ based on established criteria.^17,18^ Patients waited at least 3 months after DBS implantation to participate in the study. Exclusion criteria included neurological conditions such as: history of stroke; history of serious head injury (any neurologic sequelae, open skull fracture or hospitalization); history of definite encephalitis or oculogyric crises; drug-induced parkinsonism; sustained remission from PD; strictly unilateral features after 3 years; supranuclear gaze palsy; cerebellar signs (ataxia of gait or limbs, central nystagmus, scanning dysarthria or truncal ataxia); early severe autonomic involvement; early severe dementia (within first year of onset) with disturbances of memory, language, and praxis; extensor plantar reflex; Mini Mental Status Examination score < 24; ^16^ any defect on brain imaging (such as infarcts, brain tumor, hydrocephalus or congenital defects like lissencephaly but not cavum septum pellucidum); or MPTP exposure, for which patients were screened prior to DBS surgery. The demographics of the participants of the study are shown in **Table 1.**

**Table 1.** Demographics and clinical characteristics of 74 Parkinson disease research participants.

|  | Mean (S.D., range) |
| --- | --- |
| Age (years) | 62 (9.1, 43-80) |
| Education (years) <sup>^</sup> | 15.1 (2.7, 10-20) |
| Disease duration (years) | 12.4 (5.1, 0.51-26.5) |
| Time since STN DBS surgery (months) | 18.2 (16.1, 3-77) |
|  | **Distribution** |
| Sex | 50 male, 24 female |
| Race <sup>*</sup> | 65 Caucasian, 4 Native American/Alaskan Native, 1 African American, 1 Asian, 2 Unknown/Other |
| More affected side, by UPDRS III subscore | 41 right, 33 left |
| Dominant hand <sup>*</sup> | 65 right, 7 left, 1 ambidextrous |
| Current PD medication <sup>a, b</sup> | 74 CD-LD, 12 CD-LD ER, 33 DA agonist <sup>c</sup>, 7 MAO inhibitor, 32 COMT Inhibitor, 25 benzodiazepines, 40 amantadine, 7 antidepressants <sup>d</sup>, 21 other meds |

CD-LD = carbidopa-levodopa; CD-LD ER = carbidopa-levodopa extended release; DA = dopamine; MAO = monoamine oxidase; COMT = catechol-O-methyl transferase. <sup>^</sup>, 4 participants missing data; <sup>\*</sup>, 1 participant missing data; <sup>a</sup>, prior to abstinence on the day of study; <sup>b</sup>, participant may fall in more than one medication category; <sup>c</sup>, no participant was taking extended release formulations of DA agonists; <sup>d</sup>, amitriptyline, bupropion, duloxetine, nortriptyline, trazodone

### STN DBS Electrode Contact Selection

The side of the brain contralateral to the more affected side of the body was stimulated. The more affected side of the body was defined by the side of the body that had higher UPDRS scores in the off medication, off stimulation state.^6^ The DBS electrode contacts for each individual were placed in atlas space using a validated method^19,20^ to identify the contact locations with respect to the STN. Dorsal and ventral STN DBS contacts were chosen for each participant based on examination of their position in atlas space. Specifically, a contact within 2mm of the ventral STN border was chosen as the ventral contact, and a contact within 2mm of the dorsal STN border was chosen as the dorsal contact, ideally with one unused contact in between.^6^

### Stimulation Protocol

Participants stopped PD medications at midnight before the morning of the study. The UPDRS ratings and mood and cognitive tasks were completed during separate dorsal, ventral, and OFF STN DBS sessions over the course of one day. The order of the dorsal, ventral, and OFF sessions was randomized and blinded to the participants and raters. The voltage, frequency, and pulse width were 2.5V, 185 Hz, and 60 μs, respectively, for most participants. However, 14 participants experienced side effects from 2.5V and voltage was reduced to 1.6-2.3V.

### Measurements

Motor symptoms were rated with the Unified Parkinson Disease Rating Scale (UPDRS), part III- motor, administered by a trained clinician blind to stimulation condition. UPDRS subscale scores for bradykinesia, rigidity, tremor at rest, and total were summed contralateral to the stimulated side of the brain.

Cognition was evaluated via the spatial delayed response (SDR) and the Go/No-Go (GNG) tasks. The SDR task assesses short-term and working memory for spatial information, and was performed as described previously; the variable of interest was the distance between actual and recalled (after a 15-second delay) cue locations, or error.^21,22^ The GNG task assessed the ability to select and inhibit a pre-potent motor response appropriately under conditions of high pre- potent response strength,^23^ and was performed as described previously.^6^ The discriminability index, Pr, was the outcome measure, defined as the proportion of hits minus the proportion of false alarms. Only data from participants who reached a criterion of Pr > 0.5 in the OFF DBS condition was included in the analyses.

Current affective state was assessed using visual analog scales (VAS) and transformed to valence and arousal scores as described previously.^15,24^ Separate scores for anxiety and apathy were also measured using VAS.^8^ Higher scores on valence, anxiety and apathy represented better state.

### Primary Statistical Analyses

#### Outliers

In the data sets for all measures in both statistical analyses—univariate and 3D— outliers were defined as data values more than 3 standard deviations from the mean. The data sets and statistical outcomes shown are based on the data sets with these outliers removed.

#### Univariate statistics

To determine whether STN DBS conditions induced changes in motor, cognitive, and mood measures, we calculated difference scores by subtracting OFF condition scores from dorsal and ventral DBS condition scores to obtain “dorsal DBS difference scores” and “ventral DBS difference scores”, respectively. Dorsal and ventral DBS scores were compared using paired *t*-tests, for total contralateral UPDRS, tremor at rest, rigidity, bradykinesia, SDR error in mm, Go-No-Go Pr, valence, arousal, apathy, and anxiety.

#### Statistical mapping of DBS effects to STN anatomy

Our mapping method is described in detail in Eisenstein et al.^15^ Briefly, four statistical maps were generated for each measure. 1) An *N*-image shows the number of stimulated contacts that contributed dorsal or ventral DBS difference scores to each voxel of the map, *i.e.* within 1.3 mm. Voxels with *N* < 6 were not included in further steps. 2) A weighted mean image, containing the weighted mean difference scores across participants, with nearer contacts weighted higher. 3) A *t* image depicting weighted *t* values derived from single-sample *t* tests comparing the mean difference scores (dorsal – OFF or ventral – OFF) at each voxel to zero. 4) A *p* image containing *p* values for the *t* test at each voxel.

#### Type I error correction for multiple comparisons and sample bias

To test whether the anatomical location of the active DBS contact significantly contributed to clinical effects we used a permutation test as previously described. ^15^ Briefly, for each measure, a summary score reflecting the extent and amplitude of significant voxels in the *p* image was generated, and compared to 1000 summary scores generated similarly but from randomly chosen pairings of the active contact locations and difference scores. We considered a *p*-value ≤ 0.05 (*i.e.*, a summary score that would place it in the top 50 of the 1000 random data permutations) to indicate that DBS location significantly contributed to a measure’s difference scores.

## Results

### Distribution of Contacts

The stimulated contacts from 74 participants, each with a dorsal STN and ventral STN contact, are shown in **Figure 1**. All contacts were located within 2 mm of the STN border.

**Figure 1.**
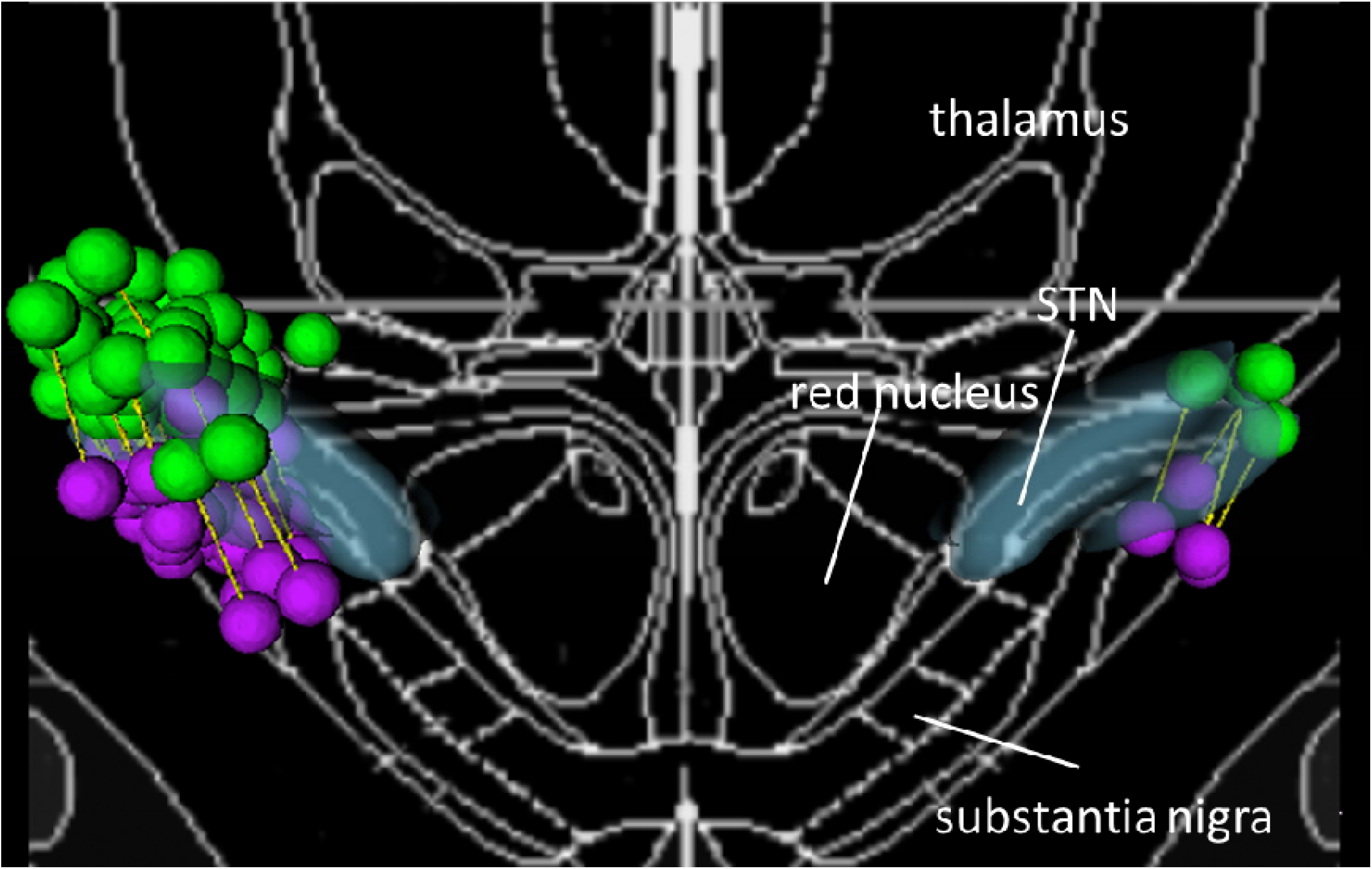
Distribution of contacts included in the analyses shown as green (dorsal) and purple (ventral) spheres, with paired contacts of each participant indicated by yellow connecting rods, and blue transparent regions indicating the subthalamic nucleus (STN).

### Univariate Results

Effects of dorsal or ventral STN DBS on mood, cognitive and motor measures (irrespective of 3D active contact location) are described in **Table 2**. Ventral or dorsal DBS significantly improved total contralateral UPDRS motor score and subscales for rigidity, tremor at rest, bradykinesia, and anxiety. Ventral DBS (only) significantly improved apathy and affective valence. Unilateral STN DBS did not significantly affect the mean scores for the Go/No-Go and SDR cognition tests. Dorsal scores differed significantly from ventral scores for anxiety, valence, and rigidity, with anxiety and valence improving more with ventral DBS, and rigidity improving more with dorsal DBS.

**Table 2.**
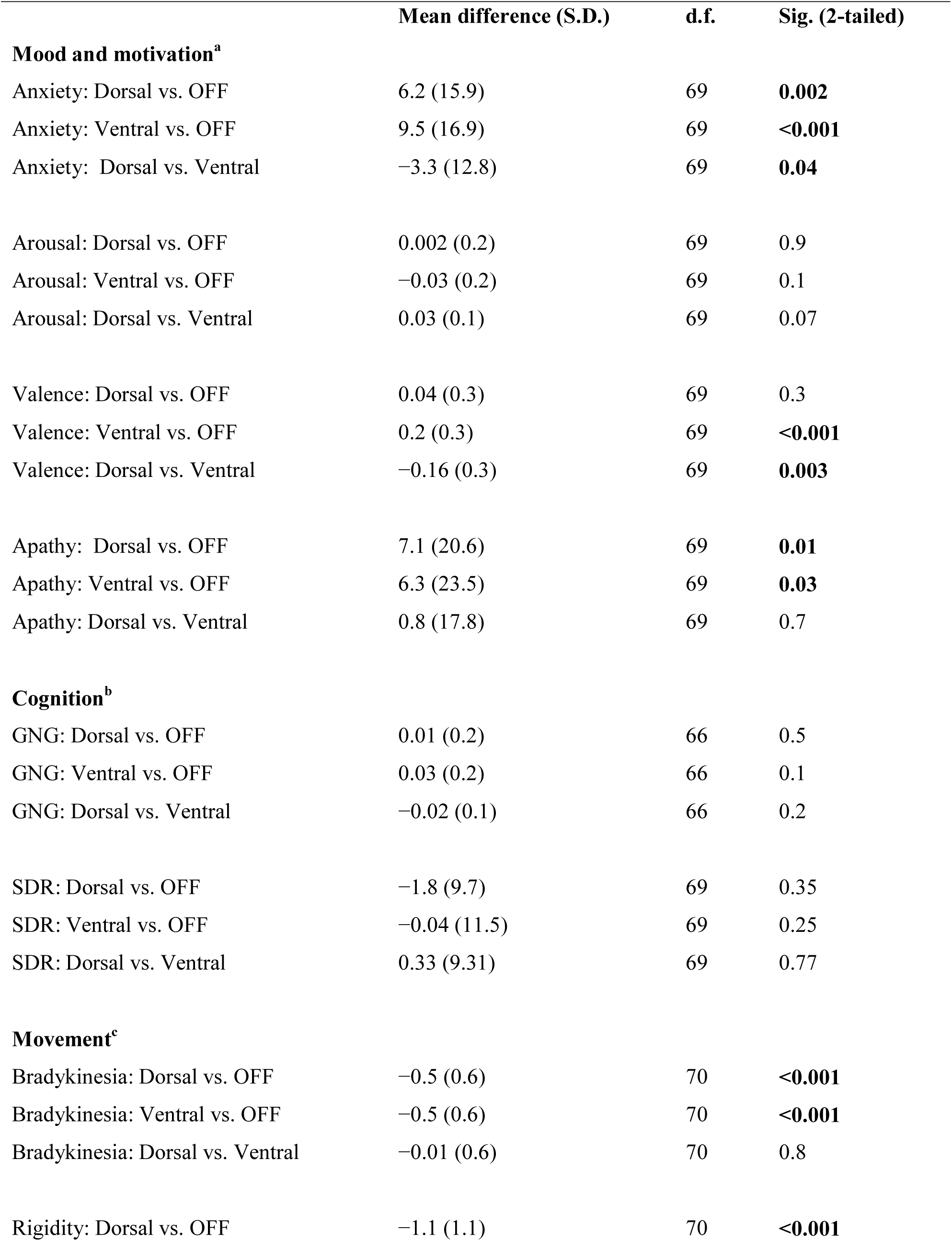
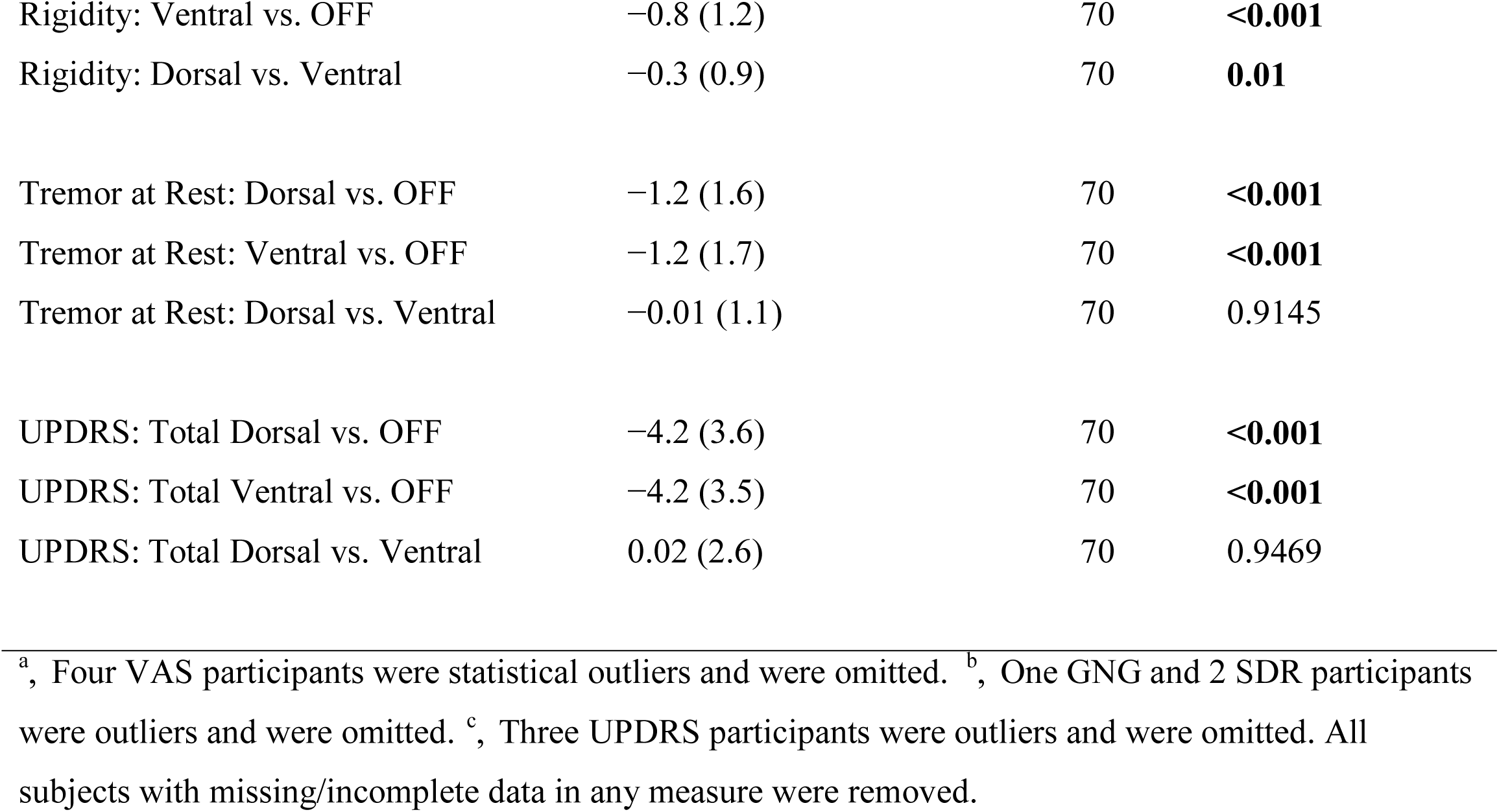
Outcome measures, by STN DBS conditions and DBS site (dorsal *vs.* ventral STN).

|  | Mean difference (S.D.) | d.f. | Sig. (2-tailed) |
| --- | --- | --- | --- |
| **Mood and motivation<sup>a</sup>** |  |  |  |
| Anxiety: Dorsal vs. OFF | 6.2 (15.9) | 69 | **0.002** |
| Anxiety: Ventral vs. OFF | 9.5 (16.9) | 69 | **<0.001** |
| Anxiety: Dorsal vs. Ventral | -3.3 (12.8) | 69 | **0.04** |
| Arousal: Dorsal vs. OFF | 0.002 (0.2) | 69 | 0.9 |
| Arousal: Ventral vs. OFF | -0.03 (0.2) | 69 | 0.1 |
| Arousal: Dorsal vs. Ventral | 0.03 (0.1) | 69 | 0.07 |
| Valence: Dorsal vs. OFF | 0.04 (0.3) | 69 | 0.3 |
| Valence: Ventral vs. OFF | 0.2 (0.3) | 69 | **<0.001** |
| Valence: Dorsal vs. Ventral | -0.16 (0.3) | 69 | **0.003** |
| Apathy: Dorsal vs. OFF | 7.1 (20.6) | 69 | **0.01** |
| Apathy: Ventral vs. OFF | 6.3 (23.5) | 69 | **0.03** |
| Apathy: Dorsal vs. Ventral | 0.8 (17.8) | 69 | 0.7 |
| **Cognition<sup>b</sup>** |  |  |  |
| GNG: Dorsal vs. OFF | 0.01 (0.2) | 66 | 0.5 |
| GNG: Ventral vs. OFF | 0.03 (0.2) | 66 | 0.1 |
| GNG: Dorsal vs. Ventral | -0.02 (0.1) | 66 | 0.2 |
| SDR: Dorsal vs. OFF | -1.8 (9.7) | 69 | 0.35 |
| SDR: Ventral vs. OFF | -0.04 (11.5) | 69 | 0.25 |
| SDR: Dorsal vs. Ventral | 0.33 (9.31) | 69 | 0.77 |
| **Movement<sup>c</sup>** |  |  |  |
| Bradykinesia: Dorsal vs. OFF | -0.5 (0.6) | 70 | **<0.001** |
| Bradykinesia: Ventral vs. OFF | -0.5 (0.6) | 70 | **<0.001** |
| Bradykinesia: Dorsal vs. Ventral | -0.01 (0.6) | 70 | 0.8 |
| Rigidity: Dorsal vs. OFF | -1.1 (1.1) | 70 | **<0.001** |
| Rigidity: Ventral vs. OFF | −0.8 (1.2) | 70 | **<0.001** |
| Rigidity: Dorsal vs. Ventral | −0.3 (0.9) | 70 | **0.01** |
| Tremor at Rest: Dorsal vs. OFF | −1.2 (1.6) | 70 | **<0.001** |
| Tremor at Rest: Ventral vs. OFF | −1.2 (1.7) | 70 | **<0.001** |
| Tremor at Rest: Dorsal vs. Ventral | −0.01 (1.1) | 70 | 0.9145 |
| UPDRS: Total Dorsal vs. OFF | −4.2 (3.6) | 70 | **<0.001** |
| UPDRS: Total Ventral vs. OFF | −4.2 (3.5) | 70 | **<0.001** |
| UPDRS: Total Dorsal vs. Ventral | 0.02 (2.6) | 70 | 0.9469 |

<sup>a</sup>, Four VAS participants were statistical outliers and were omitted. <sup>b</sup>, One GNG and 1 SDR participants were outliers and were omitted. <sup>c</sup>, Three UPDRS participants were outliers and were omitted. All subjects with missing/incomplete data in any measure were removed.

### STN DBS effects depend on DBS site

For the analysis based on 3D location of DBS, statistical significance for each measure is shown in **Table 3.** DBS location significantly contributed to the effects of STN DBS on bradykinesia and on tremor at rest. Statistical maps for these two effects are shown in **Figure 2**.

**Table 3.** Statistical summary of 3D analyses ^^^

|  | *p* | Peak<br>Weighted<br>Mean<br>Value | Peak Weighted<br>Mean Location<br>(x, y, z) | Peak<br>Weighted<br>Mean<br>Location | Peak in *p*<br>image | Peak *p* Location<br>(x, y, z) | Peak *p* Location |
| --- | --- | --- | --- | --- | --- | --- | --- |
| **Movement<sup>a</sup>** |  |  |  |  |  |  |  |
| Bradykinesia | 0.01 | -1.1 | (11.5, -14.5, -5) | cp | <0.001 | (12.9, -19.8, -4.1) | Dorsal STN |
| Rigidity | 0.30 |  |  |  |  |  |  |
| Tremor at rest | 0.02 | -4.3 | (8.5, -22.5, -4.5) | ZI | <0.001 | (13.4, -19.8, -2.9) | Dorsal STN/ZI |
| UPDRS Total | 0.08 | -9.5 | (8.5, -22.5, -6) | ZI | <0.001 | (12, -21, -3.5) | ZI |
| **Mood and motivation<sup>b</sup>** |  |  |  |  |  |  |  |
| Anxiety | 0.2 |  |  |  |  |  |  |
| Arousal | 0.2 |  |  |  |  |  |  |
| Valence | 0.1 | +0.42<br>-0.51 | (13, -23.5, -2.5)<br>(13, -13.5, 1.0) | VPM<br>H2 | <0.001<br>0.01 | (14, -19.5, -4)<br>(9.5, -16, -2.5) | Dorsal STN/cp<br>STN |
| Apathy | 0.2 |  |  |  |  |  |  |
| **Cognition<sup>c</sup>** |  |  |  |  |  |  |  |
| GNG | 0.9 |  |  |  |  |  |  |
| SDR | 0.6 |  |  |  |  |  |  |

<sup>^</sup>, Peak *p* and weighted mean values and locations are only listed for the measures found to be significant in the permutation analysis. <sup>a</sup>, Seventy-one participants contributed to the analyses for the motor measures. <sup>b</sup>, Seventy participants contributed to the analyses for the mood measures. <sup>c</sup>, Sixty-seven and 69 participants contributed to the GNG and SDR analyses, respectively. STN, subthalamic nucleus; cp, cerebral peduncle; ZI, zona incerta; VPM, ventral posterior medial thalamic nucleus; H2, lenticular fasciculus (field H2); GNG, Go-NoGo; SDR, spatial delayed response. Both positive and negative suprathreshold clusters were observed for valence difference score, so peak coordinates and weighted mean values are given for both.

**Figure 2 (see next page).**
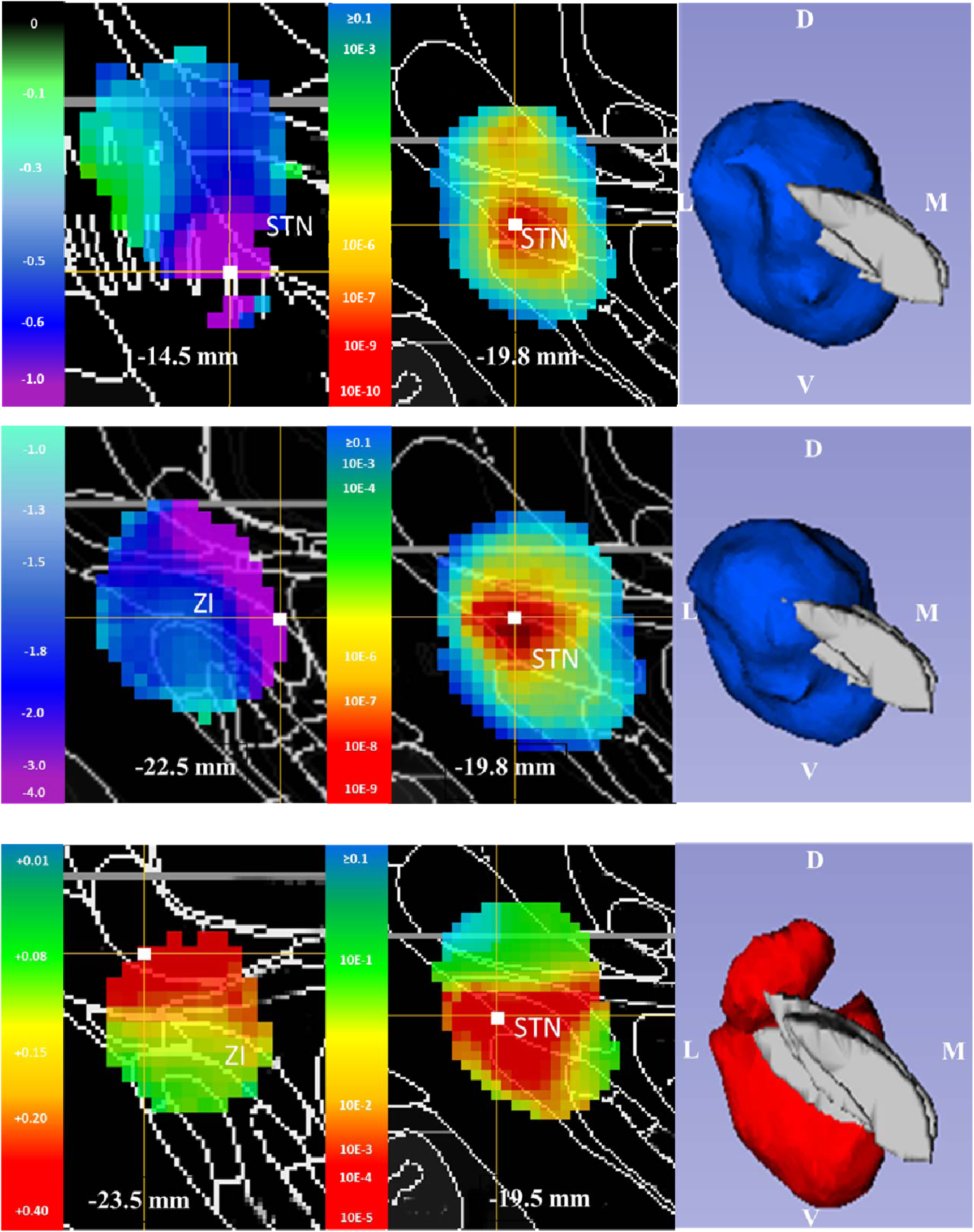
Weighted mean image, *p* image, and 3D *p* image for measures with significant effect of contact location in the 3D analyses. Upper panel: bradykinesia. Middle panel: tremor at rest. Lower panel: valence. For the weighted mean images, the cooler shades indicate where, on average, the difference scores (ventral-OFF and dorsal-OFF) are more negative (improvement relative to OFF for motor measures; worsening relative to OFF for valence), while the warmer shades indicate locations where the difference scores are more positive (worsening relative to OFF for motor measures; improvement relative to OFF for valence). For the 2D *p-*image, warmer shades indicate more significant *p* values, while the cooler shades indicate less significant *p* values. White squares indicate peak coordinates. The 3D image is shown as viewed from anteriorly, and the blue or red volume indicates values <0.05 in the *p* image (for valence, the positive cluster is shown). STN, subthalamic nucleus; ZI, zona incerta, D, dorsal, V, ventral, L, lateral, M, medial.

## Discussion

The results support the conclusion that 3D electrode contact location contributes to the motor effects of STN DBS. The peak *p* values for DBS-induced improvements in motor function were located more dorsally in the STN. This confirms our findings in a different sample, using a different experimental design,^15^ which showed greater motor improvement in dorsolateral STN, particularly for tremor at rest. Similarly, previous studies also suggested greater improvement in motor function in dorsal STN and the zona incerta (ZI).^10, 25, 26^ These results fit with anatomical data placing the dorsolateral portion of the STN in a loop connecting primary motor cortex to putamen and motor thalamus, and linking the zona incerta to motor and limbic systems.

In the current study, electrode contact site, as a 3D variable, did not significantly alter the effect of STN DBS on cognitive or mood function in PD. The lack of effect on mood may be only a Type II error: *p* was 0.1 for the affective valence permutation analysis, and ventral STN stimulation improved valence more than dorsal STN stimulation in the univariate analysis. A previous study^27^ showed increased mood improvement with STN DBS in those with anxiety or mood disorders or higher symptom severity, but psychiatric diagnosis was not assessed in the present study. The nonsignificant association of contact location and cognitive function is more surprising, given the present sample size and our previous findings that DBS effects on cognitive measures were location dependent.^6,15^ However, there are several differences between the current study and the most comparable previous study.^15^ First, the previous study’s ON sessions tested participants with their individually optimized DBS settings, including choice of active contact. In other words, in that study the contact selection was not chosen blind to clinical response.

Furthermore, since in that study the contacts and settings were optimized clinically, cognitive or affective responses may have contributed to selecting contact or pulse settings that were more likely to improve mood or thinking than the anatomically chosen DBS contacts and standardized pulse settings in the present study.

Strengths of this study include its relatively large sample size, acute stimulation paradigm, assessment blind to the location stimulated, and innovative statistical approach. Limitations include the fact that clinical DBS electrode implantation targets the dorsal posterolateral STN, which necessarily limits the number of contacts that fall in the anterior or medial-ventral STN. The limited number of data points in these regions reflects this reality, reducing power in parts of the ventromedial and anterior STN. Second, in some conditions participants or examiners may have detected when STN was turned on. However, as the focus of the study was the correlation of effect with DBS site, and neither the participants nor the examiners knew the precise locations of the contacts, the study was still blinded for the key variable under investigation, *i.e.*, the location of the active contact. Third, the minimum interval between DBS changes (42 minutes) was chosen based on previous experience with motor signs, but there may be longer-term effects—on mood and cognitive function in particular—that this investigation may have missed. However, the time limit on the OFF session was also chosen with ethical and practical considerations in mind that preclude extending the time to study more delayed effects. Lastly, statistically significant changes in rating scales may not imply syndromal or clinically significant changes.

Our previous study in a different PD sample did not support complete functional segregation within the STN of mood, motor, and cognitive function.^15^ This new sample provides some functional evidence for a dorsal–ventral, motor–non-motor gradient of benefit. Rigidity, resting tremor, and bradykinesia improved significantly more with dorsal stimulation (Table 2), and the 3D analysis found significant location effects for bradykinesia and tremor at rest (Table 3), with the evidence for improvement stronger in ZI or dorsolateral STN for both measures (Figure 2, upper panel, *p* image, and lower panel, weighted mean image). By contrast, anxiety and affective valence improved significantly more with ventral than dorsal stimulation (Table 2), and in the 3D analysis, affective valence tended to improve most with ventral STN stimulation (Figure 2, weighted mean image; *p*=0.1). On the other hand, stimulation of either ventral or dorsal STN improved motor function, anxiety and apathy, and cognitive effects did not differ with stimulation site. Therefore the direct, functional evidence supports only a mild dorsal–ventral gradient for motor and non-motor effects of STN DBS, rather than a strict dorsal–ventral functional segregation.

## Ethics

The study was approved by the Human Research Protection Office at WUSM and was carried out in accordance with the principles expressed in the Declaration of Helsinki. All participants provided written, informed consent.

## Data Availability

The data for this study appear as electronic supplementary material.

## Competing interests

We have no competing interests.

## Authors’ contributions

AG participated in data analysis and in drafting and revising the manuscript. SAE contributed to study design, data analysis, and drafting and revising the manuscript. NTT participated in data analysis and manuscript revision. JMK participated in data analysis. MCC, JSP and TH contributed to study design, data collection and manuscript revision. MU collected data and revised the manuscript. KJB contributed to study design, drafting and manuscript revision. All authors approved the final manuscript.

## Funding

National Institutes of Health (NIH; R01 NS041509 and R01 NS075321 to JSP, R01 NS058797 to TH); Michael J. Fox Foundation for Parkinson’s Research to KJB; American Parkinson Disease Association (APDA) Greater St. Louis chapter, APDA Advanced Research Center at Washington University and Barnes-Jewish Hospital Foundation (Elliot Stein Family Fund and Parkinson Disease Research Fund) to JSP, Brain & Behavior Research Foundation (NARSAD) Young Investigator Award to MCC; American Brain Foundation / American Academy of Neurology Clinical Research Training Fellowship to MU. Research reported in this publication was supported by the Washington University Institute of Clinical and Translational Sciences grant UL1TR000448 from the National Center for Advancing Translational Sciences (NCATS) of the NIH. The content is solely the responsibility of the authors and does not necessarily represent the official view of the funding agencies.

